# Phytochemical-induced mucin accumulation in the gastrointestinal lumen is independent of the microbiota

**DOI:** 10.1101/2022.03.11.483917

**Authors:** Andrew J. Forgie, Tingting Ju, Stephanie L. Tollenaar, Benjamin P. Willing

## Abstract

The mucus layer is critical to gastrointestinal health and ecology. Dietary phytochemicals are well documented to stimulate mucus production and secretion, but the underlying mechanism and effects on gut health are poorly understood. We fed germ-free and conventional mice diets containing approximately 0.4% of polyphenols per gram to determine if the phytochemical-induced accumulation of mucin in the gastrointestinal lumen is dependent on the microbiota. In addition, we assess how increased mucin shapes microbial communities in conventional mice. Germ-free mice receiving a pea (*Pisum sativuum*) seed coat proanthocyanidin-containing diet (PA) had greater levels of fecal mucin compared to the non-proanthocyanidin-containing (NPA) pea seed coat diet control (*P* < 0.05), confirming that fecal mucin accumulation is independent of the gut microbiota. Conventional mice fed the PA diet and a red osier dogwood (ROD; *Cornus sericea*) extract diet (DW) had higher mucin levels compared to a control diet without phytochemicals (*P* < 0.01 and *P* < 0.05, respectively). The increase in luminal mucin was associated with consistent increases in bacterial taxa belonging to *Lachnospiraceae* and [*Clostridium*] *leptum* species and a decrease in *Romboutsia* species. We conclude that phytochemicals have the ability to alter gut microbial ecology by increasing the amount of mucin in the gastrointestinal lumen.

## Introduction

The mucus layer of the mammalian gastrointestinal (GI) tract is a major part of the innate immune system. It is made of heavily glycosylated mucin glycoproteins produced by goblet cells and provides lubrication, hydration, and protection against pathogens and harmful substances that pass through the intestinal environment (1). Two distinctive mucus layers exist: an inner layer devoid of bacteria and an outer loose layer filled with mucolytic and associated microbes that inhabit that particular niche (2–5). Diet has been well recognized to alter host-microbe interactions in the GI tract that affect the integrity of the mucus layer and intestine (6). In particular, phytochemical consumption modulates mucus production, metabolism and has antimicrobial activities that directly affects the gut microbiota; however, the mechanism of their beneficial health outcomes is poorly understood.

Polyphenolic compounds make up the majority of the bioactive phytochemicals consumed in the human diet (7). They are categorized as hydrolysable or condensed (non-hydrolysable) high molecular weight tannins (gallotannins & ellagitannins, and proanthocyanidins, respectively) and low molecular weight polyphenols that include phenolic acids, flavonoids, lignans, stilbenes, and curcumins (8). Because condensed tannins are resistant to host acid and enzyme hydrolysis, they are more likely to reach the gut microbiota compared to the quickly absorbed hydrolysable tannins and low molecular weight phytochemicals. Health benefits have been attributed to their free radical scavenging capacity to neutralize inflammation-causing reactive oxygen species, as well as their direct antimicrobial effect on microbial communities, and the indirect production of bioactive polyphenolic catabolites by the gut microbiota (9,10).

Phytochemicals have an ability to alter mucus physiology, and along with their numerous forms and bioactivities they have shown contradictory effects on resisting pathogen colonization and virulence in the GI tract (11,12). Phytochemical research has focused on improving intestinal barrier integrity with the mechanism of action hinting towards its ability to stimulate the mucus layer, with an increase in mucus production and thickness considered beneficial (11,13,14). However, thickness and expression of mucus-related genes does not necessarily mean that the mucus layer has formed properly for protection. In agreement, a previous study conducted by our group examining the supplementation of peas (Pisum sativuum) rich or low in polyphenol content showed that the polyphenol-rich pea diet increased the amount of fecal mucin in the lumen (12). Excess mucin in the GI lumen was associated with greater Citrobacter rodentium colonization and activation of a proinflammatory response. This diet-induced mucus phenotype has been experimentally tested and confirmed in vitro, as indicated by the ability of galloylated tannins and related compounds to directly cross-link with purified mucins, thereby altering the viscoelastic properties of mucus (15). Therefore, phytochemical consumption increases the accessibility of mucus glycans to the gut microbiota, but to what extent this drives gut ecology has not been determined.

In this study, we investigated whether the presence of microbes is required for the previously observed increase in fecal mucin in response to polyphenol-rich pea seed coat consumption. Whether changes in the gut microbiota drives the mucus phenotype or is a consequence of host-diet interactions remains unknown. We hypothesized that luminal mucin accumulation in the GI tract from phytochemical supplementation is dependent on the microbiota. We fed the proanthocyanidin (PA) and non-proanthocyanidin (NPA) high (20% w/w)-fat pea diets used in our previous study to germ-free (GF) mice and measured fecal mucin (12). In addition, we tested how the non-hydrolysable PA diet compares to a hydrolysable red-osier dogwood (ROD; Cornus sericea) extract on fecal mucin and microbial communities when fed to conventional mice. ROD extracts have been shown in pig models to improve feed efficiency and promote gut resistance to invading pathogens (16); however, the underlying mechanism is poorly understood and possibly driven by changes to the mucus layer. The identification of mucin-degrading bacteria and their impact on the gut niche environment in response to dietary phytochemicals will help determine their contribution to gut ecology and health.

## Material and methods

### Animals and dietary treatments

All animal experiments were conducted in accordance with guidelines set by the Canadian Council on Animal Care and approved by the Animal Care and Use Committee at the University of Alberta (Edmonton, AB, Canada). All mice used in this study were bred and maintained in the University of Alberta Axenic Mouse Research Unit. Mice were eight to ten-weeks-old and allowed to acclimatize on an autoclaved standard chow diet (5010 maintenance diet, LabDiet, St. Louis, MO, USA) for a week prior to dietary treatments. Table 1 provides formulations of the isocaloric treatment diets, which were balanced for macronutrients and insoluble fiber with cellulose. Eight female GF Swiss-Webster mice were housed four per open-top cage in the same GF isolator (CEP Standard Safety, McHenry, Illinois, USA) and handling was done directly inside the isolator. GF mice were fed treatment diets containing pea seed coats flours rich (‘Solido’ cultivar; PA) and poor (‘Canstar’ cultivar; NPA) in proanthocyanidin content as describe previously (12). The diets provided to GF mice were prepared with 15 g instead of 10 g of vitamin mix per kg diet to account for the loss imposed by irradiating the diets to 10 kGY, which was done at the Cross-Cancer Institute at the University of Alberta. Germ free status following the diet intervention was confirmed by anaerobic and aerobic culture of fecal pellets at termination. To investigate the role of microbes and test a second polyphenolic source on the fecal mucin phenotype, we fed a control diet (Control), a PA diet (PA), and the ROD supplemented diet (DW) to conventional Swiss-Webster mice (Table 1). Mice were housed two to four per cage using the Tecniplast Isocage-P bioexclusion system (Buguggiate, VA, Italy) and all animal handling was done in a biosafety cabinet. Control and PA diet groups included two male and two female mice housed separately to determine if sex plays role in the fecal mucin phenotype, previously identified in female mice only (12). The spray-dried ROD extract (Red Dog Enterprises Ltd., Winnipeg, MB, Canada) was added to the dogwood (DW) diet at 4% as done in previous studies (16). ROD extracts can contain bioactive phenolic compounds at 4% to 22% depending on the season (17). The addition of polyphenolic extracts was calculated based on the average total phenolic content of 10% in the ROD extract for the DW diet and 4.51% in the pea seed coat flour for the PA diet. The final amount of proanthocyanidin content in the diets was 0.4% per gram of diet. All diets were prepared aseptically as powdered diet at the University of Alberta and mice had ad libitum access to water and diet throughout the two-week long diet intervention. Body weights were taken every second day and fecal samples were collected aseptically at beginning (day 0) and end (day 14) of the dietary treatment for conventional mice. Mice were euthanized using carbon dioxide. Samples were collected aseptically and stored at −80°C until use. previously and adjusted accordingly. Red osier dogwood extract was provided by Roberts Scales of Red Dog Enterprises Ltd. Diets are isocaloric and contains 0.4% (w/w) of total polyphenols (pea seed coat flour = 4.51%; red osier dogwood extract = 10%).

**Table 1.** Composition of high (20% w/w)-fat dietary treatments (g/kg).

| Component, g/kg | Germ-free |  | Conventional |  |  |
| --- | --- | --- | --- | --- | --- |
|  | NPA | PA | Control | PA | DW |
| Lard (Tenderflake) | 190 | 190 | 190 | 190 | 190 |
| Flaxseed oil | 2 | 2 | 2 | 2 | 2 |
| Corn oil (Mazola) | 8 | 8 | 8 | 8 | 8 |
| Casein | 267 | 262 | 270 | 262 | 267 |
| L-Methionine | 2.5 | 2.5 | 2.5 | 2.5 | 2.5 |
| Dextrose | 214 | 214 | 214 | 196 | 195 |
| Corn Starch | 194 | 194 | 193.4 | 180.7 | 180.4 |
| Cellulose | 0 | 0 | 50 | 0 | 45 |
| Mineral Mix | 51 | 51 | 51 | 51 | 51 |
| Vitamin Mix | 15 | 15 | 10 | 10 | 10 |
| Inositol | 6.3 | 6.3 | 6.3 | 6.3 | 6.3 |
| Choline Chloride | 2.8 | 2.8 | 2.8 | 2.8 | 2.8 |
| Canstar (NPA) seed coat | 71.5 |  |  |  |  |
| Solido (PA) seed coat |  | 96.5 |  | 88.7 |  |
| Red osier dogwood (DW) |  |  |  |  | 40 |
| **Total Weight (g)** | **1024.1** | **1044.1** | **1000** | **1000** | **1000** |
| Fat / g | 0.2 | 0.2 | 0.2 | 0.2 | 0.2 |
| Protein / g | 0.3 | 0.3 | 0.3 | 0.3 | 0.3 |
| Carbohydrate / g | 0.4 | 0.4 | 0.4 | 0.4 | 0.4 |
| Insoluble fiber / g | 0.04 | 0.05 | 0.05 | 0.05 | 0.05 |
| Energy (At Water) kcal / g | 4.4 | 4.3 | 4.5 | 4.4 | 4.4 |
| **% PACs / g** | **0** | **0.4** | **0** | **0.4** | **0.4** |

Note: The nutrient content of NPA ('canstar') and PA ('solido') cultivars were analyzed previously and adjusted accordingly. Red osier dogwood extract was provided by Roberts Scales of Red Dog Enterprises Ltd. Diets are isocaloric and contains 0.4% (w/w) of total polyphenols (pea seed coat flour = 4.51%; red osier dogwood extract = 10%).

### Fecal mucin assay

Fecal pellets from individual mice were pooled across daily fecal collections throughout the last four days of the two-week dietary treatment. Pooled fecal collections were subsequently freeze-dried and ground to a powder. A fluorometric assay kit (Fecal Mucin Assay kit; Cosmo Bio co. LTD, Carlsbad, CA, USA) that quantifies N-acetylgalactosamine, the reducing end sugar of the O-linked glycan chain, was used to determine the mucus content (18).

### Microbial community analyses

Total DNA was extracted from colon contents using the QIamp Fast DNA Stool Mini Kit (Qiagen, Valencia, CA, USA) with an additional bead-beating step using ~200 mg of garnet rock at 6.0 m/s for 60 s on a FastPrep-24 5G instrument (MP Biomedicals, Irvine, CA, USA). Paired-end sequencing was accomplished using the Illumina MiSeq Platform (2 x 300 cycles; Illumina Inc., San Diego, CA, USA). Amplicon libraries were constructed according to the protocol from Illumina (16S Metagenomic Sequencing Library Preparation) that amplified the V3-V4 region of the bacterial 16S rRNA gene: 341F (5’ - TCGTCGGCAGCGTCAGATGTGTATAAGAGACAGCCTACGGGNGGCWGCAG-3’) and 805R (5’ - GTCTCGTGGGCTCGGAGATGTGTATAAGAGACAGGACTACHVGGGTATCTAATCC-3’). Raw sequences were processed with Quantitative Insight into Microbial Ecology 2 (QIIME 2) (19) pipeline using DADA2 to filter, trim and merge paired-end reads into amplicon sequence variants (ASVs). Phylogenetic trees were constructed using the qiime alignment (mafft; mask) and qiime phylogeny (fasttree; midpoint-root) function. Taxonomy was assigned using the qiime feature-classifier classify-sklearn function using the SILVA v138 database trained for the specific amplicon region (20). QIIME2 files (.qza) were imported into R using qiime2R (version 0.99.4) package and analyzed with phyloseq (version 1.34.0) package (21). Sequences belonging to ‘chloroplast’ and ‘mitochondria’ were removed. In addition, ‘Lactococcus’ sequences were dropped from the analysis because they were suspected to be a contaminant from casein (22). Numbers assigned to ASVs reflect their total counts from highest to lowest count across samples. Alpha diversity (Observed, Shannon, phylogenetic diversity (PD)) and beta diversity based on a Bray-Curtis dissimilarity index were done with rarefied reads at a count of 8444.

### Statistical analysis

Significance testing and graphing for body weights and fecal mucin were done in GraphPad Prism 6 (Graphpad Software, LaJolla, CA. USA) using a t-test or a parametric anova corrected for multiple comparisons with Tukey’s post-hoc test. Data are presented as mean ± standard error of the mean. Letters were used to denote a significance when appropriate. Statistical significance for alpha diversity was determine with the anova TukeyHSD() correction function. Principal coordinate analyses (PCoA) was plotted using the phyloseq package and clustering significance was determined using the ‘betadisper’ function (23) for dispersion and ‘pairwiseAdonis.dm’ function (24) for orientation. Differential abundance analysis was done with DESeq2 using non-rarefied reads and tree_glom() function to merge similar ASVs. The ‘log2foldchange’ of only the ASVs with a P value less than 0.05 were plotted with bolded ASVs signifying the significant adjusted P value < 0.10, < 0.05 (*), < 0.01 (**) and < 0.001 (***).

## Results

### Phenolic compounds increase mucin content in the gut independently of the microbiota

GF mice fed the PA diet had higher amounts of fecal mucin (P < 0.05) compared to the NPA control diet (Fig 1a). Phytochemicals directly increased fecal mucin in the GI tract independently of the microbiota. The presence of microbes did not alter this outcome as conventional mice displayed a similar increase in fecal mucin in the PA (P < 0.05), as well as the DW (P < 0.05) diet compared to control (Fig 1b). Polyphenolic diets did not affect the weights of conventional mice over the course of the two-week experiment (Fig 1c). Conventional female and male mice fed the Control and PA diets displayed no difference in weight gain or amount of fecal mucin. Although the limited sample size may not adequately represent the sex effect, the impact of phytochemicals on fecal mucin was not different between sexes.

**Fig 1.**
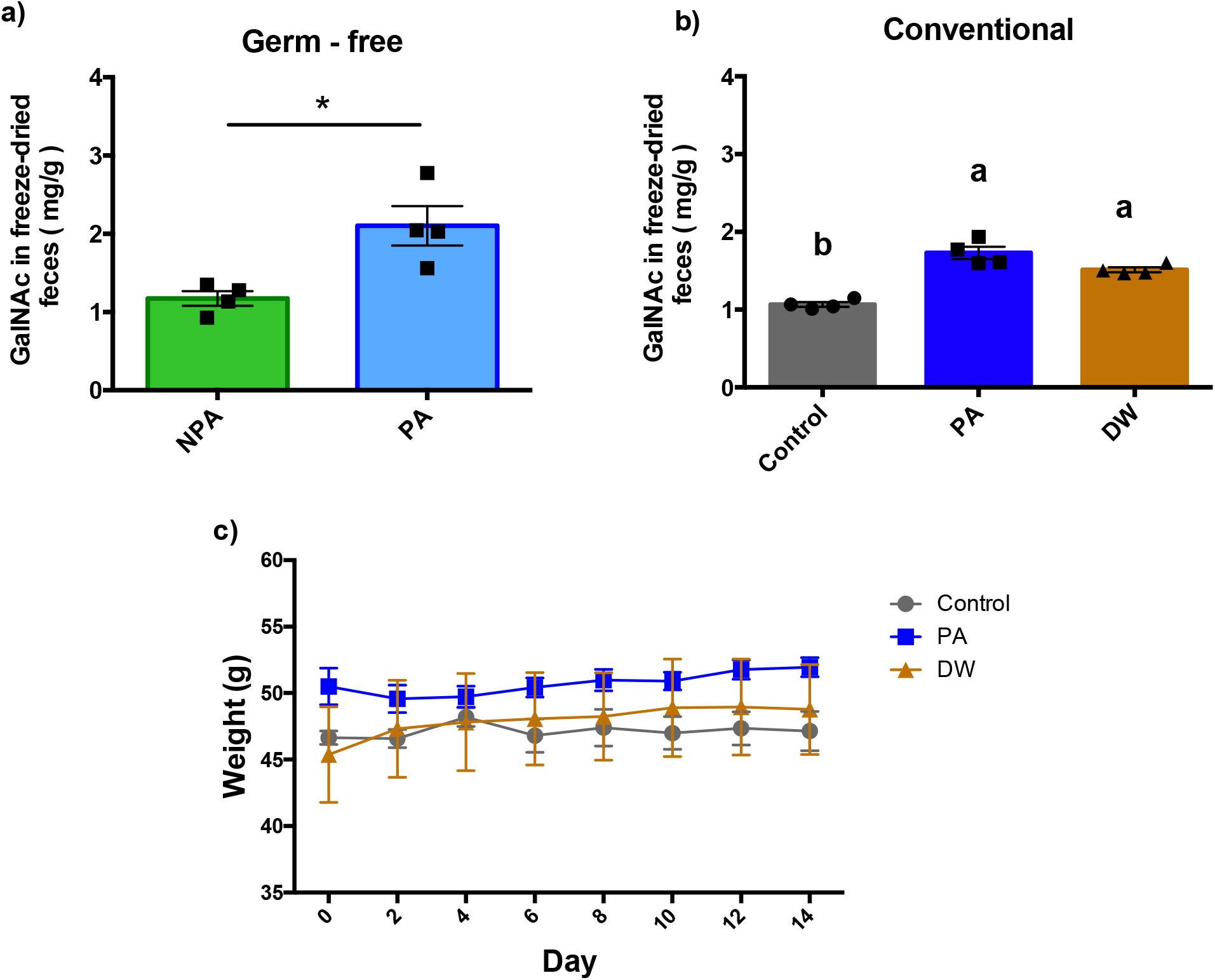
Phytochemical diets (PA and DW) increased the mucin content in the gastrointestinal tract independently of the microbiota. (a) Germ-free Swiss-Webster mice fed the PA diet responded by increasing fecal mucin content, a novel finding that suggest phytochemicals act directly on host mucus chemistry independently of the microbiota (n = 4; * *P* < 0.05). (b) Dietary phytochemicals significantly increased fecal mucin content in PA (*P* < 0.01) and DW (*P* < 0.05) groups of conventional mice (n = 4). (c) Conventional Swiss-Webster mice weights were unaffected by dietary treatment (n = 4).

### Changes in microbial composition in response to phenolic compound rich diets

Phytochemical diets associated with the fecal mucin phenotype revealed similar changes to microbial communities. PCoA was conducted using Bray-Curtis dissimilarity metric to visualize microbial communities before and after dietary treatment and identify overall differences between treatments. We analyzed microbial communities without separating female and male mice because sex did not appear to alter the amount of mucin recovered in fecal samples. Microbial communities prior to diet intervention were not different between groups (Adonis unadjusted day 0 to control; PA: R2 = 0.10, P = 0.67 & DW: R2 = 0.11; P = 0.56) but clustered distinctly after 14 days (Adonis unadjusted day 14 to control; PA: R2 = 0.27, P = 0.11 & DW: R2 = 0.79, P = 0.02). All dietary treatments led to distinct changes in microbial communities when compared to their initial microbial community (Adonis unadjusted day 0 to day 14; Control: R2 = 0.53, P = 0.05; PA: R2 = 0.55, P = 0.04 & DW: R2 = 0.76, P = 0.05) (Fig 2a). Dispersion analysis between groups did not pass significance and shows that the within treatment variability was consistent between groups. PCoA of day 14-microbial communities revealed that both PA and DW diets had similar changes to microbial communities but were still distinct from one another. Principal component (PC) 1, PC2 and PC3 explains 68.2%, 10.3%, and 8.8% respectively and when visualized as PC1 vs PC2 (Fig 2b) and PC2 vs PC3 (Fig 2c) distinct clustering between treatments can be visualized. Alpha diversity metrics of day 0 and day 14 microbial communities show that all dietary treatments reduced the unique counts (Observed; P < 0.05); however, diversity as determined by Shannon index revealed that all but the DW group (P < 0.05) remained constant compared to both Control and PA groups (Fig 2d). The PD index revealed that Control and PA diets had lower microbial diversity (P < 0.05) at day 14 compared to day 0, whereas the DW group maintained a similar diversity as at day 0 before diet treatment (Fig 2d).

**Fig 2.**
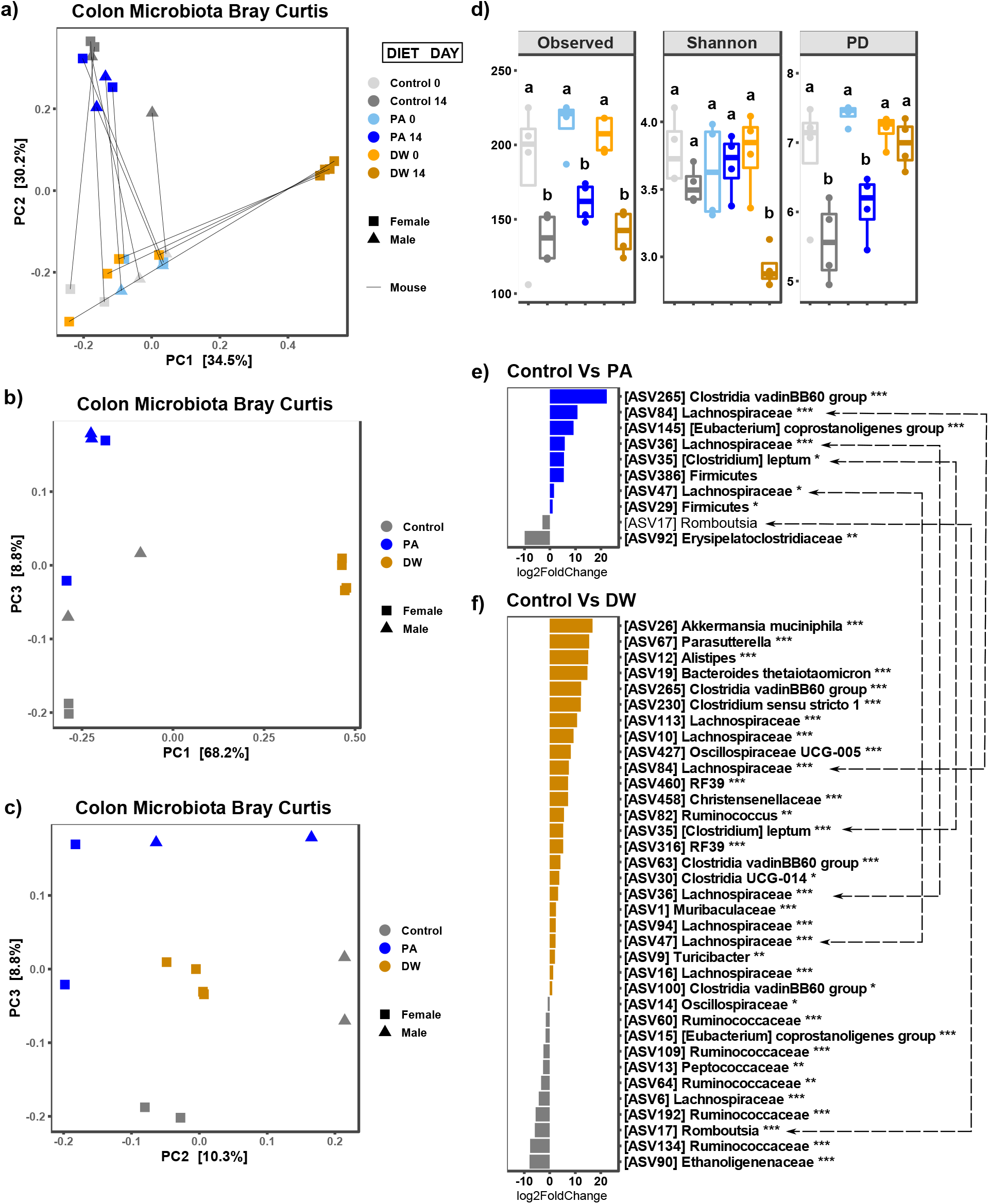
Phytochemical diets had a distinct impact on the microbial community structure with consistent changes to *Lachnospiraceae, Clostridium*, and *Romboutsia* species in the colon of conventional mice. (a) Microbial communities prior to diet intervention were not different between groups (Bray-Curtis PCoA; Adonis unadjusted day 0 to control; PA: R2 = 0.10, P = 0.67 & DW: R2 = 0.11, P = 0.56) but clustered distinctly after 14 days (Bray-Curtis PCoA; Adonis unadjusted day 14 to control; PA: R2 = 0.27, P = 0.11 & DW: R2 = 0.79, P = 0.02). All mice had similar microbial communities Control: R2 = 0.53, P = 0.04; PA: R2 = 0.55, P = 0.05 & DW: R2 = 0.76, P = 0.03).

The phytochemical diets drastically altered the colonic microbiota as determined by differential expression of ASVs using DESeq2 compared to control (Fig 2e-f). This includes numerous ASVs assigned to taxa belonging to the Firmicutes phylum. Compared to the control group, the PA diet reduced *Romboutsia* and *Erysipelatoclostridiaceae*, while increasing *Lachnospiraceae*, [*Clostridium*] *leptum*, [*Eubacterium*] *coprostanoligenes*, and a bacterium from the *Clostridia* vadinBB60 group (Fig 2e). The DW diet reduced *Romboutsia*, *Ruminococcaceae* members, *Oscillospiraceae*, [*Eubacterium*] *coprostanoligenes*, *Peptococcaceae*, and *Ethanoligenenaceae* members of the Firmicute population (Fig 2f). The DW diet increased *Akkermansia muciniphila*, *Parasutterella*, *Alistipes*, *Turicibacter*, and *Bacteroides thetaiotaomicron*, along with an unclassified member from the *Muribaculaceae* family.

At day 14, diet (b) Principal coordinate analysis plot of day 14 microbial communities using principal component 1 (PC1; 68.2%) and principal component 3 (PC3; 8.8%), along with (c) principal component 2 (PC2; 10.3%) plotted against PC3 shows a distinct but subtle similarity between dietary groups. (d) Alpha diversity metrics (Observed, Shannon, PD) of day 0 and day 14 microbial communities shows that all treatment diets reduced diversity (P < 0.05); however, DW diet specifically reduced diversity (Observed and Shannon; P < 0.05) compared to both Control and PA groups. Differential expression of ASVs as determine by DESeq2 were plotted for (e) PA and (f) DW compared to Control group. Consistent ASVs that respond to PA and DW diets are noted with hashed arrow lines, this includes an increase in Lachnospiraceae and [Clostridium] leptum species along with a decrease in a Romboutsia species (n = 4; bolded taxa represent a trend (P < 0.10); * P < 0.05, ** P < 0.01, *** P < 0.001).

## Discussion

Phytochemicals are secondary metabolites produced by plants to communicate with their environment. They are generally considered non-essential nutrients that contribute to physiology through numerous bioactive properties ranging from antioxidant, anti-inflammatory, and antimicrobial to protein chelation and enzyme inhibition (25). Thousands of phytochemicals have been identified and are noted for their free radical scavenging capacity to neutralize reactive oxygen species associated with inflammation and disease (10). Polymeric phytochemicals, such as proanthocyanidins, are the most abundant and bioactive polyphenols in our diet (26). They are composed of polymeric flanva-3-ol monomers, known as condensed tannins and linked through double A-type and single B-type linkages (26). The degree and type of polymerization, along with the galloylated moieties determine their physicochemical structure and thus their bioactivities on both the gut microbiota and host (27,28). The bioactive properties of polyphenolic compounds and their impact on physiology have made them good candidates for antibiotic alternatives and therapeutics against enteric infection (29).

In this study, we determined that the increase in fecal mucin in response to polyphenol-rich diets observed previously (12,30) occurs independently of the gut microbiota. The inclusion of polyphenols in diet at 0.4% in this study appears safe, as no change in body weight was documented throughout the two-week dietary intervention. This is in accordance with previous studies that show that similar amounts of phytochemicals in diets do not negatively impact growth, and may even improve it (12,16). Despite being quite different with regards to phenolic compound composition, the PA and DW diets both led to increased fecal mucin, suggesting that flavan-3-ol condense tannins from peas, beans or fruit are not unique in their ability to increase fecal mucin (12,30,31). This occurs independently of the gut microbiota; however, more studies are required to determine the main phytochemicals or groups of chemicals responsible for the mucin phenotype. Our results provide *in vivo* support to the previous *in vitro* experiments showing that phytochemicals, particularly the galloylated polyphenols, directly disrupt the viscoelastic properties of mucus by disrupting binding among mucin glycoproteins (15). In our previous study (12), the pea seed coat supplementation (PA diet) led to faster colonization of *C. rodentium*, a common enteric mouse pathogen, and we suspect that this is a consequence the direct impact of polyphenols on the mucus layer. Most studies have focused on the beneficial roles that phytochemicals have on health and has been extensively reviewed (32); however, little is mentioned of the phytochemical-mucus interactions in the gut outlined herein. Therefore, in addition to confirming that phytochemical directly increase luminal mucin concentration, an analysis of the colonic microbial community was conducted, with a particular interest in mucolytic microbes that may benefit under these conditions.

Microbial community analysis revealed that pea seed coat and ROD-supplementation alters the gut microbiota. The PCoA plot analyses revealed that the DW diet substantially altered microbial composition compared to the PA diet. Shannon diversity and PD values of the colonic microbiome in the PA group is consistent with our previous experiment (12). The reduced Shannon diversity of the DW group could be explained by the antimicrobial properties of ROD phytochemicals; however, more research is required to confirm the direct antimicrobial actions of ROD supplementation. A study in pigs with a lower inclusion rate of 0.5% ROD extract showed no effect on ileal microbial alpha diversity (Shannon and Simpson) but a prebiotic effect on Lactobacillus species was noted along with no change to growth performance (33). A study in weaned pigs challenged with *Escherichia coli* k88+ found that 2% and 4% ROD extract diets conferred beneficial effects on growth performance; however, microbial composition was not assessed (16). Phylogenetic diversity in the DW group was maintained at day 14, which could be explained by the increased abundance of *Akkermansia municiphila*, *Parasutterella* and *Turicibacter,* which only appeared in this DW group at day 14 and were not detected in any group at day 0. Although we did not detect these microbes in the sequencing data of Control and PA groups at day 14, they may have been present below our detection limit for 16s rRNA sequencing. The ROD extract effectively reduced the abundance of some species thereby opening up a niche for others. For this reason, we see higher PD values in the DW group compared to Control and PA groups.

Microbes that were enriched by both phenolic diets include *Lachnospiraceae* and [*Clostridium*] *leptum* species, which may reflect a response to increased mucin availability. Luminal mucin levels were confirmed greater in both PA and DW diet groups compared to Control. We characterized an increase in abundance of *A. muciniphila*, a well-known mucolytic microbe, in the DW group but not the PA group. Mucin supplementation has been confirmed in mice to encourage mucin degrading bacteria, such as *A. muciniphila*, and mitigates diet-induced microbiota perturbations (34). Moreover, phytochemicals are well-known to increase the abundance of *A. muciniphila* and improved health outcomes (35). The phytochemical-induced mucin phenotype may partly explain their increased abundance in the gut; however, the absence of *A. muciniphila* in the PA diet suggests other factors contributed to their increase in the DW group. A study in mice using jaboticaba fruit, which is high in flavan-3-ols, at 5%, 10%, and 15% of diet found an increase in *A. muciniphila* at only 10% and 15% (36) suggesting a dose-dependent threshold that supports their growth likely exists in the gut. The lack of *A. muciniphila* in the PA group of this present study is inconsistent with our previous study (12) and suggest mice did not receive the necessary dose of polyphenols to encourage *A. muciniphila* fitness in the gut. The moderate effect of the PA diet on microbial communities could be explained a loss in the antimicrobial actions of pea phytochemicals after long-term storage. As a result, the PA diet had less severe impacts on microbial community structure as compared to the DW diet and suggested that the interactions among microbes are stable enough to prevent *A. muciniphila* expansion even with increased access to mucin glycans. In addition, both PA and DW diets led to a consistent decrease in abundance of *Romboutsia*, a species that is highly adapted to nutrient-rich environments (37) and a potential marker of stability in the gut. The competitive advantage gained by mucolytic bacteria may have altered the nutrient-rich niches in the gut that genera like *Romboutsia* depend on for growth. The mucus layer supports microbial niches by directly providing glycans for energy and indirectly through cross-feeding from one microbe to another (5). Future research should focus on characterizing the bioactive compounds promoting luminal mucin accumulation, as well as identifying the type and source of the accumulating mucins (1). Further knowledge of these interactions will provide the foundational framework necessary to understand how the mucus layer contributes to host-microbe stability and health.

## Conclusion

The production and maintenance of the mucus layer is a vital part of intestinal homeostasis. It has become clear that host mucus provides a foundation of host derived glycans that supports mutualism and commensalism among microbes in the gut. Understanding how phytochemicals influence the viscoelasticity of the mucus layer will help to determine the best therapeutic use of phytochemicals to promote health. This research provides insight into establishing the mechanisms involved in the ability of mucus to stabilize gut ecology and control microbial communities. Further studies are required to determine the specific phytochemical compounds and structure that induce mucus secretion and/or disrupt mucin binding and formation.

## Acknowledgments

We would like to thank Roberts Scales and Dr. Jocelyn Ozga for providing red osier dogwood extract and pea seed coats respectively.

## References

1. Dhanisha SS, Guruvayoorappan C, Drishya S, Abeesh P. Mucins: Structural diversity, biosynthesis, its role in pathogenesis and as possible therapeutic targets. Crit Rev Oncol Hematol [Internet]. 2018;122(December 2017):98–122. Available from: https://doi.org/10.1016/j.critrevonc.2017.12.006

2. Johansson MEV, Jakobsson HE, Holmén-Larsson J, Schütte A, Ermund A, Rodríguez-Piñeiro AM, et al. Normalization of Host Intestinal Mucus Layers Requires Long-Term Microbial Colonization. Cell Host Microbe [Internet]. 2015 Nov;18(5):582–92. Available from: https://linkinghub.elsevier.com/retrieve/pii/S1931312815004175

3. Jakobsson HE, Rodriguez-Pineiro AM, Schutte A, Ermund A, Boysen P, Bemark M, et al. The composition of the gut microbiota shapes the colon mucus barrier. EMBO Rep [Internet]. 2015;16(2):164–77. Available from: http://embor.embopress.org/cgi/doi/10.15252/embr.201439263

4. Paone P, Cani PD. Mucus barrier, mucins and gut microbiota: the expected slimy partners? Gut [Internet]. 2020 Dec;69(12):2232–43. Available from: https://gut.bmj.com/lookup/doi/10.1136/gutjnl-2020-322260

5. Li H, Limenitakis JP, Fuhrer T, Geuking MB, Lawson MA, Wyss M, et al. The outer mucus layer hosts a distinct intestinal microbial niche. Nat Commun [Internet]. 2015;6(May):8292. Available from: http://www.nature.com/doifinder/10.1038/ncomms9292

6. Forgie AJ, Fouhse JM, Willing BP. Diet-Microbe-Host Interactions That Affect Gut Mucosal Integrity and Infection Resistance. Front Immunol [Internet]. 2019 Aug 6;10(August):1802. Available from: http://www.ncbi.nlm.nih.gov/pubmed/31447837

7. Pandey KB, Rizvi SI. Plant polyphenols as dietary antioxidants in human health and disease. Oxid Med Cell Longev [Internet]. 2009;2(5):270–8. Available from: http://www.ncbi.nlm.nih.gov/pubmed/20716914%5Cnhttp://www.pubmedcentral.nih.gov/articlerender.fcgi?artid=PMC2835915

8. Crozier A, Jaganath IB, Clifford MN. Phenols, Polyphenols and Tannins: An Overview. In: Crozier A, Clifford MN, Ashihara H, editors. Plant Secondary Metabolites [Internet]. Oxford, UK: Blackwell Publishing Ltd; 2006. p. 1–24. Available from: http://doi.wiley.com/10.1002/9780470988558

9. Kawabata K, Yoshioka Y, Terao J. Role of Intestinal Microbiota in the Bioavailability and Physiological Functions of Dietary Polyphenols. Molecules [Internet]. 2019 Jan 21;24(2):370. Available from: http://www.mdpi.com/1420-3049/24/2/370

10. Li A-N, Li S, Zhang Y-J, Xu X-R, Chen Y-M, Li H-B. Resources and Biological Activities of Natural Polyphenols. Nutrients [Internet]. 2014 Dec 22;6(12):6020–47. Available from: http://www.mdpi.com/2072-6643/6/12/6020

11. Wlodarska M, Willing BP, Bravo DM, Finlay BB. Phytonutrient diet supplementation promotes beneficial Clostridia species and intestinal mucus secretion resulting in protection against enteric infection. Sci Rep [Internet]. 2015;5:9253. Available from: http://www.pubmedcentral.nih.gov/articlerender.fcgi?artid=4365398&tool=pmcentrez&rendertype=abstract

12. Forgie AJ, Gao Y, Ju T, Pepin DM, Yang K, Gänzle MG, et al. Pea polyphenolics and hydrolysis processing alter microbial community structure and early pathogen colonization in mice. J Nutr Biochem [Internet]. 2019 May 8;67:101–10. Available from: http://www.ncbi.nlm.nih.gov/pubmed/30877891

13. Bergstrom KSB, Kissoon-Singh V, Gibson DL, Ma C, Montero M, Sham HP, et al. Muc2 Protects against Lethal Infectious Colitis by Disassociating Pathogenic and Commensal Bacteria from the Colonic Mucosa. Roy CR, editor. PLoS Pathog [Internet]. 2010 May 13;6(5):e1000902. Available from: https://dx.plos.org/10.1371/journal.ppat.1000902

14. Arike L, Hansson GC. The Densely O-Glycosylated MUC2 Mucin Protects the Intestine and Provides Food for the Commensal Bacteria. J Mol Biol [Internet]. 2016 Aug 14;428(16):3221–9. Available from: https://linkinghub.elsevier.com/retrieve/pii/S0022283616001157

15. Georgiades P, Pudney PDA, Rogers S, Thornton DJ, Waigh TA. Tea Derived Galloylated Polyphenols Cross-Link Purified Gastrointestinal Mucins. Haverkamp RG, editor. PLoS One [Internet]. 2014 Aug 27;9(8):e105302. Available from: https://dx.plos.org/10.1371/journal.pone.0105302

16. Koo B, Amarakoon SB, Jayaraman B, Siow YL, Prashar S, Shang Y, et al. Effects of dietary red-osier dogwood (Cornus stolonifera) on growth performance, blood profile, ileal morphology, and oxidative status in weaned pigs challenged with Escherichia coli K88 +. Miglior F, editor. Can J Anim Sci [Internet]. 2021 Mar 1;101(1):96–105. Available from: https://cdnsciencepub.com/doi/10.1139/cjas-2019-0188

17. Isaak CK, Petkau JC, Karmin O, Ominski K, Rodriguez-Lecompte JC, Siow YL. Seasonal variations in phenolic compounds and antioxidant capacity of Cornus stolonifera plant material: Applications in agriculture. Can J Plant Sci [Internet]. 2013 Jul;93(4):725–34. Available from: http://www.nrcresearchpress.com/doi/10.4141/cjps2012-310

18. Crowther RS, Wetmore RF. Fluorometric assay of O-linked glycoproteins by reaction with 2-cyanoacetamide. Anal Biochem [Internet]. 1987 May;163(1):170–4. Available from: https://linkinghub.elsevier.com/retrieve/pii/0003269787901084

19. Bolyen E, Rideout JR, Dillon MR, Bokulich NA, Abnet CC, Al-Ghalith GA, et al. Reproducible, interactive, scalable and extensible microbiome data science using QIIME 2. Nat Biotechnol [Internet]. 2019 Aug 24;37(8):852–7. Available from: http://www.nature.com/articles/s41587-019-0209-9

20. Bokulich NA, Kaehler BD, Rideout JR, Dillon M, Bolyen E, Knight R, et al. Optimizing taxonomic classification of marker-gene amplicon sequences with QIIME 2’s q2-feature-classifier plugin. Microbiome [Internet]. 2018 Dec 17;6(1):90. Available from: https://microbiomejournal.biomedcentral.com/articles/10.1186/s40168-018-0470-z

21. McMurdie PJ, Holmes S. phyloseq: an R package for reproducible interactive analysis and graphics of microbiome census data. PLoS One [Internet]. 2013;8(4):e61217. Available from: http://www.ncbi.nlm.nih.gov/pubmed/23630581

22. Kimoto H, Nomura M, Kobayashi M, Mizumachi K, Okamoto T. Survival of lactococci during passage through mouse digestive tract. Can J Microbiol [Internet]. 2003 Nov 1;49(11):707–11. Available from: http://www.nrcresearchpress.com/doi/10.1139/w03-092

23. Anderson MJ. Distance-based tests for homogeneity of multivariate dispersions. Biometrics [Internet]. 2006 Mar;62(1):245–53. Available from: http://www.ncbi.nlm.nih.gov/pubmed/16542252

24. Martinez Arbizu P. pairwiseAdonis: Pairwise multilevel comparison using adonis. 2017.

25. Balaji M, Ganjayi MS, Hanuma Kumar GEN, Parim BN, Mopuri R, Dasari S. A review on possible therapeutic targets to contain obesity: The role of phytochemicals. Obes Res Clin Pract [Internet]. 2016 Jul;10(4):363–80. Available from: http://dx.doi.org/10.1016/j.orcp.2015.12.004

26. Monagas M, Urpi-Sarda M, Sánchez-Patán F, Llorach R, Garrido I, Gómez-Cordovés C, et al. Insights into the metabolism and microbial biotransformation of dietary flavan-3-ols and the bioactivity of their metabolites. Food Funct [Internet]. 2010;1(3):233. Available from: http://xlink.rsc.org/?DOI=c0fo00132e

27. Yokota K, Kimura H, Ogawa S, Akihiro T. Analysis of A-Type and B-Type Highly Polymeric Proanthocyanidins and Their Biological Activities as Nutraceuticals. J Chem [Internet]. 2013;2013:1–7. Available from: http://www.hindawi.com/journals/jchem/2013/352042/

28. Girard M, Bee G. Invited review: Tannins as a potential alternative to antibiotics to prevent coliform diarrhea in weaned pigs. Animal [Internet]. 2020;14(1):95–107. Available from: http://dx.doi.org/10.1017/S1751731119002143

29. Willing BP, Pepin DM, Marcolla CS, Forgie AJ, Diether NE, Bourrie BCT. Bacterial resistance to antibiotic alternatives: a wolf in sheep’s clothing?1. Anim Front [Internet]. 2018 Jun 7;8(2):39–47. Available from: https://academic.oup.com/af/article/8/2/39/4990302

30. Taira T, Yamaguchi S, Takahashi A, Okazaki Y, Yamaguchi A, Sakaguchi H, et al. Dietary polyphenols increase fecal mucin and immunoglobulin A and ameliorate the disturbance in gut microbiota caused by a high fat diet. J Clin Biochem Nutr [Internet]. 2015 Nov;57(3):212–6. Available from: https://www.jstage.jst.go.jp/article/jcbn/57/3/57_15-15/_article

31. Tanaka M, Honda Y, Miwa S, Akahori R, Matsumoto K. Comparison of the Effects of Roasted and Boiled Red Kidney Beans (Phaseolus vulgaris L.) on Glucose/Lipid Metabolism and Intestinal Immunity in a High‐Fat Diet‐Induced Murine Obesity Model. J Food Sci [Internet]. 2019 May 16;84(5):1180–7. Available from: https://onlinelibrary.wiley.com/doi/10.1111/1750-3841.14583

32. Andersen-Civil AIS, Arora P, Williams AR. Regulation of Enteric Infection and Immunity by Dietary Proanthocyanidins. Front Immunol [Internet]. 2021 Feb 24;12(February):1–12. Available from: https://www.frontiersin.org/articles/10.3389/fimmu.2021.637603/full

33. Zheng S, Song J, Qin X, Yang K, Liu M, Yang C, et al. Dietary supplementation of red-osier dogwood polyphenol extract changes the ileal microbiota structure and increases Lactobacillus in a pig model. AMB Express [Internet]. 2021 Dec 29;11(1):145. Available from: https://doi.org/10.1186/s13568-021-01303-8

34. Pruss KM, Marcobal A, Southwick AM, Dahan D, Smits SA, Ferreyra JA, et al. Mucin-derived O-glycans supplemented to diet mitigate diverse microbiota perturbations. ISME J [Internet]. 2021;15(2):577–91. Available from: http://dx.doi.org/10.1038/s41396-020-00798-6

35. Anhê FF, Pilon G, Roy D, Desjardins Y, Levy E, Marette A. Triggering Akkermansia with dietary polyphenols: A new weapon to combat the metabolic syndrome? Gut Microbes [Internet]. 2016;7(2):146–53. Available from: http://www.ncbi.nlm.nih.gov/pubmed/26900906%5Cnhttp://www.pubmedcentral.nih.gov/articlerender.fcgi?artid=PMC4856456

36. Soares E, Soares AC, Trindade PL, Monteiro EB, Martins FF, Forgie AJ, et al. Jaboticaba (Myrciaria jaboticaba) powder consumption improves the metabolic profile and regulates gut microbiome composition in high-fat diet-fed mice. Biomed Pharmacother [Internet]. 2021 Dec;144:112314. Available from: https://linkinghub.elsevier.com/retrieve/pii/S0753332221010982

37. Gerritsen J, Hornung B, Renckens B, van Hijum SAFT, Martins dos Santos VAP, Rijkers GT, et al. Genomic and functional analysis of Romboutsia ilealis CRIB T reveals adaptation to the small intestine. PeerJ [Internet]. 2017 Sep 11;5(9):e3698. Available from: https://peerj.com/articles/3698

